# Ludicrous Speed Linear Mixed Models for Genome-Wide Association Studies

**DOI:** 10.1101/154682

**Authors:** Carl Kadie, David Heckerman

**Affiliations:** Microsoft Research Redmond, WA 98052

## Abstract

We have developed Ludicrous Speed Linear Mixed Models, a version of FaST-LMM optimized for the cloud. The approach can perform a genome-wide association analysis on a dataset of one million SNPs across one million individuals at a cost of about 868 CPU days with an elapsed time on the order of two weeks. A Python implementation is available at https://fastlmm.github.io/.

**Significance:** Identifying SNP-phenotype correlations using GWAS is difficult because effect sizes are so small for common, complex diseases. To address this issue, institutions are creating extremely large cohorts with sample sizes on the order of one million. Unfortunately, such cohorts are likely to contain confounding factors such as population structure and family/cryptic relatedness. The linear mixed model (LMM) can often correct for such confounding factors, but is too slow to use even with algebraic speedups known as FaST-LMM. We present a cloud implementation of FaST-LMM, called Ludicrous Speed LMM, that can process one million samples and one million test SNPs in a reasonable amount of time and at a reasonable cost.

## Introduction

Identifying SNP-phenotype correlations using genome-wide association studies (GWAS) is difficult because effect sizes are so small for common, complex diseases. To address this issue, institutions are creating extremely large cohorts with sample sizes on the order of one million. Unfortunately, such cohorts are likely to contain confounding factors such as population structure and family/cryptic relatedness, which leads to inflated type-I errors when analyzed with traditional methods.

The linear mixed model (LMM) can often correct for such confounding factors [1]. Unfortunately, in its original form, its computational complexity of runtime and memory made it prohibitively expensive to use. Relatively recently, improvements through algebraic transformations known as FaST-LMM, have made it possible to scale LMM computations to sample sizes of about 100 thousand [2,3].

Here, we present a cloud implementation of FaST-LMM, called Ludicrous Speed LMM. Ludicrous Speed LMM can process one million samples and one million test SNPs in a reasonable amount of time, at a reasonable cost, and with arbitrarily little memory provided extremes (e.g., testing one SNP at a time or all SNPs at once) are avoided.

## Methods

We begin with a description of linear mixed models and FaST-LMM. The basic idea behind the linear mixed model is that a single test SNP is regressed on a trait, with *K* other SNPs acting as covariates. For reasons that will become clear shortly, we will refer to these covariates as similarity SNPs. Let *y_i_*, *s_i_*, and **G***_i_* = (*g*_*i*1_, …, *g_iK_*) denote the trait, test SNP, and *K* similarity SNPs for the *i*th individual, respectively. Let **y** = (*y*_1_, …, *y_N_*)^T^, **s** = (*s*_1_, …, *s_N_*)^T^, and 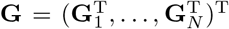 denote the observations of the trait, test SNP, and *K* similarity SNPs, respectively, across the individuals. Thus, **G** is an *N* × *K* matrix, where the *ij*th element corresponds to the *j*th similarity SNP of the *i*th individual. We model the influence of the test SNP and similarity SNPs on the trait as follows:

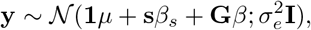

where *μ* is an offset and **1** is column of ones, *β_s_* is the weight relating the test SNP to the trait, *β*^T^ = (*β*_1_, …, *β_K_*) are the weights relating the similarity SNPs to the trait, 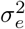 is a scalar, and 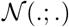 denotes the multivariate normal distribution.

Taking a Bayesian approach, we assume that each of the *β_i_* corresponding to the similarity SNPs are mutually independent, each having a normal distribution with the same variance

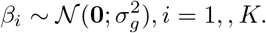

Further, we standardize the observations of each similarity SNP across the individuals to have variance 1 (and mean 0) so that, *a priori*, each SNP has an equal influence on the trait. We similarly standardize the test SNP.

Averaging over the distributions of the *β_i_*, we obtain

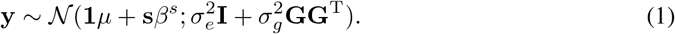

The distribution in (1) is a linear mixed model [1,2]. As we have just shown, it corresponds to a Bayesian linear regression, also known as L2-regularized linear regression. The distribution also corresponds to a Gaussian process with a linear covariance or kernel function [4]. The model implies that the correlation between the traits of two individuals is related to the dot product of the similarity SNPs for those two individuals, hence the name similarity SNPs. The similarity matrix **GG**^T^ is known as the *Realized Relationship Matrix* (RRM) [5]. In general, other similarity measures can be and have been used [5].

From the perspective of the linear mixed model as a Bayesian linear regression, it is clear that the test and similarity SNPs should be disjoint. Otherwise, we would be conditioning on the SNP we are trying to test. Moreover, due to linkage disequilibrium, we should avoid the use of similarity SNPs that are near the test SNP. Doing otherwise has been termed *proximal contamination* [2]. In practice, when computing *P* values for test SNPs on a given chromosome, **G** is typically built with similarity SNPs from all but that chromosome [2]. We employ this practice here. Also, when the elements of **G** are scaled so that its diagonal sums to *N* (the expected value of the diagonal), the estimate of narrow-sense heritability is more accurate [7]. Our implementation of Ludicrous Speed LMM includes this scaling.

An F-test [6] is used to compute a *P* value for each test SNP. First, the parameters of the model (*μ, β_s_, σ_e_, σ_g_*) are fit with restricted maximum likelihood (REML) for the null and alternative hypotheses. The parameters can be computed in closed form except the ratio of 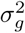 to 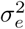, which is usually (and herein) determined via grid search [2]. This ratio is computed for the null hypothesis and also applied to the alternative hypothesis [2].

As we mentioned in the introduction, a straightforward implementation of GWAS based on (1) is computationally inefficient. Namely, *P*-value computations require manipulations of **GG**^T^ that scale cubically with sample size *N*, yielding an overall runtime complexity of *O*(*MN*^3^) when testing *M* SNPs. Thus, the model is infeasible for GWAS with sample sizes greater than 10^4^.

The FaST-LMM algorithm employs algebraic transformations, allowing computations to scale to sample sizes on the order of 10^5^. FaST-LMM consists of two key transformations. First, if we factor **GG**^T^ into the matrix product **UDU**^T^, where **U** is an orthogonal matrix and **D** is a diagonal matrix (a procedure known as spectral decomposition), then it can be shown that (1) can be re-written as the linear regression

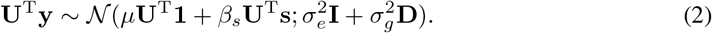

That is, the model takes the form of a linear regression after the data is rotated by **U**^T^. GWAS based on (2) can be used to test *M* SNPs with a runtime complexity of *O*(*MN*^2^). The second transformation makes use of the fact that when an RRM is used for the similarity matrix, its spectral decomposition can be replaced by an SVD of **G**. (With **G** = **U**Σ**V**^T^, **G***G*^T^ = **U**Σ^2^**U**^T^.) Furthermore, when the number of similarity SNPs *K* is less than the sample size *N*, the SVD of **G** can be replaced by a skinny **SVD** of **G** (**G**^T^**G** = **V**Σ^2^**V**^T^ yields **V** and Σ. **G** = **U**Σ**V**^T^ yields **U** by matrix multiplication.) The resulting model can be used to test *M* SNPs with a runtime complexity of *O*(*MNK*).

The condition *K* < *N* can often be satisfied, because linkage disequilibrium allows us to build **G** from a subset of available SNPs while still maintaining control of type-I error. In practice, *K* should be chosen so that there is no visible inflation in the resulting quantile-quantile QQ plots of actual P values versus expected P values under the null hypothesis. A practical approach to identifying a suitable value for *K* is to start with a small value, and then increase it until no inflation is observed. SNPs should be selected such that any two adjacent SNPs are roughly equally correlated.

There is one important remark that should be made before we move to a description of our improvements. From the Bayesian-linear-regression formulation of the linear mixed model, it is clear that the test and similarity SNPs should be disjoint. Otherwise, we would be conditioning on the SNP we are trying to test. Moreover, due to linkage disequilibrium, we should avoid the use of similarity SNPs that are near the test SNP. Doing otherwise has been termed *proximal contamination* [2]. In practice, when testing SNPs on a given chromosome, **G** is typically built with similarity SNPs from all but that chromosome [2]. We employ this practice here.

## Improvements to FaST-LMM: Ludicrous Speed LMM

Here, we describe Ludicrous Speed LMM, a cloud implementation of FaST-LMM including improvements of parallelization, block decomposition, and multithreading. We describe the improvements across the stages of analysis, partitioned as follows. For concreteness, we assume that we are analyzing human autosomal chromosomes with test SNPs appearing in each of the 22 chromosomes.

- Stage 0, **G**: Read **G**_0_ (the pre-standardized similarity SNPS), standardize them, regress out any covariates, and output **G**.
- Stage 1, **G**^T^**G**: Compute **G**^T^**G**.
- Stage 2, SVD: For each chromosome, remove the entries of **G**^T^**G** corresponding to the chromosome, and compute the singular value decomposition (SVD) on the remaining product.
- Stage 3, PostSVD: For each removal, compute the corresponding rotation matrix **U**, and identify the optimal ratio of 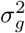 to 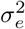.
- Stage 4, TestSNPs: For each test SNP, read its data, standardize the SNP, regress out covariates, use the appropriate **U**, and compute the P value for the SNP.

We optimized stage 0 by (1) reading selected SNPs in batches to keep the memory use arbitrarily small, (2) reading and standardizing SNPs on multiple processes, and (3) computing the sum of squares across individuals for each similarity SNP. The result of step 3 is a length *K* vector, used to scale **G** in the SVD step (see below). Calculating the vector here, but using it later, allows us to create just a single **G**, instead of needing to create 22 **G** matrices, one for each chromosome removal. We write **G** to disk as a two-dimensional array of doubles. It can be accessed via memory mapping or by streaming in blocks. In later stages, we will see how it can be used without loading all of it in memory.

We optimized stage 1, the computation of **G**^T^**G**, by (1) distributing the calculation to compute it in blocks, (2) using a tree copy to put the whole **G** file on each compute node on a solid-state drive (SSD), (3) tree scaling, that is allocating compute nodes only when there is a source for them to tree copy from, and deallocating when there is no more work for a node to do, (4) using sub-blocks for the computation of each block (allowing arbitrarily little memory to be used), and (5) doing the local calculations via multithreaded C++ (with one thread reading from the SSD and the others multiplying, yielding a CPU bounded procedure). Tree scaling reduces our compute costs with only minor effects on the elapsed compute time.

**Table 1.** CPU use per stage.

| Stage | CPU days |
| --- | --- |
| Stage 0, $\mathbf{G}$ | 1 |
| Stage 1, $\mathbf{G}^T \mathbf{G}$ | 12 |
| Stage 2, SVD | 9 |
| Stage 3, PostSVD | 19 |
| Stage 4, TestSNPs | 827 |
| Total | 868 |

We optimized stage 2, the computation of the SVDs, by (1) distributing the computations across 22 compute nodes, one for each chromosome, (2) computing the SVDs using LAPACK’s divide-and-conquer algorithm, and (3) after computing the SVD, using the sum of squares vector created in stage 0 to adjust the results to match the scaling we would have obtained had we operated on the scaled **G** matrix. Regarding step 2, the LAPACK algorithm scales as *N*^2.8^, and MKL provides an optimized, multithreaded version of it. The default MKL version doesn’t work because its integer indexes are too small, but the MKL ILP64 version works well.

We optimized stage 3 by (1) using tree copy and tree scaling, now to get **G** on each compute node on an SSD, (2) accessing **G** in blocks to make memory use arbitrarily small, and (3) using multithreaded matrix multiplication for high CPU utilization.

The computations in stage 4 are dominated by the multiplication of the test SNPs by **U**^T^. We optimized this multiplication by (1) dividing the test SNPs into blocks and distributing the work for each block across compute nodes, (2) keeping each **U** file in separate cluster storage so that all 22 files can be downloaded to their first compute node with little interference with the others, using tree copy and tree scaling for each chromosome so that each compute node needs only one of the 22 **U** files (each such file is large, on the order of 400 GB), (3) using sub-blocks to avoid large memory use as before, and (4) doing the local calculations via multithreaded C++ so that the calculations are CPU bound. Note that each compute node needs only a small portion of the test SNPs and so downloads only that small portion.

## Generation of data for testing

As we did not have access to data from a large cohort for testing, we generated synthetic data. One million SNPs were generated across one million samples with an allele-frequency distribution taken from human data. The SNPs were assigned to chromosomes in proportion to human DNA. Traits were generated at random with mean 2/3 and a standard deviation of 3. Two covariates were generated at random, each with mean 1.5 and a standard deviation of 2.

## Results

We applied Ludicrous Speed LMM on the synthetic data set using up to 115 compute nodes on an Azure cluster with D15v2 compute nodes (20 processors each). 50,000 similarity SNPs were used.

Total cluster storage was about 10 TB. The largest memory use on a single node was 140 GB. Total computation time (not counting node startup and monitoring) was 868 CPU days. Table 1 shows CPU use per stage. Figure 1 shows the CPU use per task. Generally, the cost of each chromosome is proportional to its size. The exceptions were caused by failures requiring partial restarts. In terms of elapsed time, the run took 19 days, but would have taken 9 days with no restarts. If 1000 nodes had been used without restarts, the run would have taken 5 days, and the CPU cost would have increased only 9% due to copying large files to more machines.

**Figure 1:**
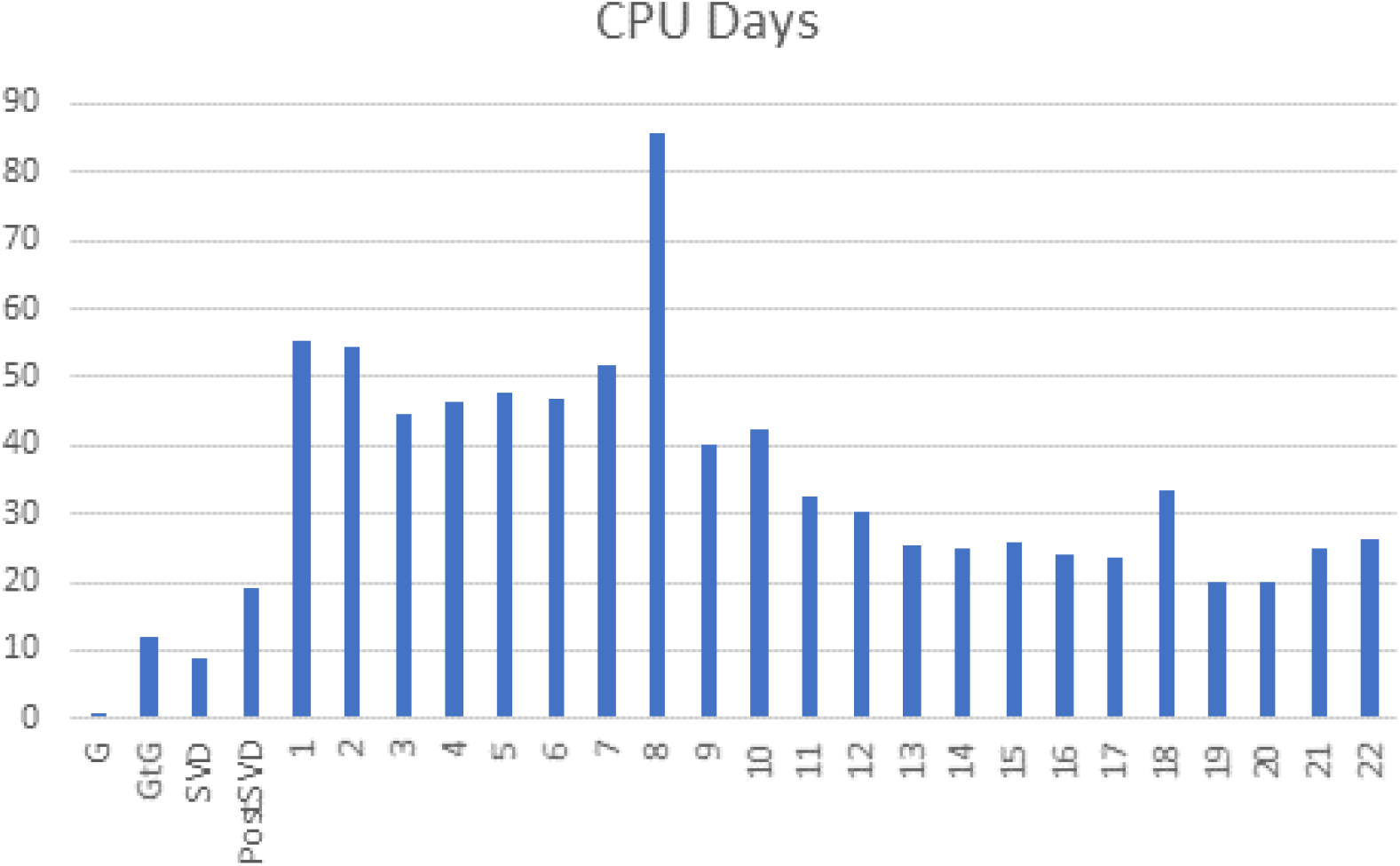
CPU use per task.

## Summary

We have developed Ludicrous Speed LMM, a version of FaST-LMM optimized for the cloud. Using 50,000 similarity SNPs, the approach can analyze a dataset of one million test SNPs across one million individuals at a cost of about 868 CPU days with an elapsed time on the order of two weeks.

A Python implementation of Ludicrous Speed LMM is available at https://fastlmm.github.io/.

